# DrivR-Base: A Feature Extraction Toolkit For Variant Effect Prediction Model Construction

**DOI:** 10.1101/2024.01.16.575859

**Authors:** Amy Francis, Colin Campbell, Tom Gaunt

## Abstract

**Motivation:** Recent advancements in sequencing technologies have led to the discovery of numerous variants in the human genome. However, understanding their precise roles in diseases remains challenging due to their complex functional mechanisms. Various methodologies have emerged to predict the pathogenic significance of these genetic variants. Typically, these methods employ an integrative approach, leveraging diverse data sources that provide critical insights into genomic function. Despite the abundance of publicly available data sources and databases, the process of navigating, extracting, and pre-processing features for machine learning models can be daunting. Furthermore, researchers often invest substantial effort in feature extraction, only to later discover that these features lack informativeness.

**Results:** In this paper, we present *DrivR-Base*, an innovative resource that efficiently extracts and integrates molecular information (features) for single nucleotide variants from a wide range of databases and tools, including AlphaFold, ENCODE, and *Variant Effect Predictor*. The resulting features can be used as input for machine learning models designed to predict the pathogenic impact of human genome variants in disease. Moreover, these feature sets have applications beyond this, including haploinsufficiency prediction and the development of drug repurposing tools. We describe the resource’s development, practical applications, and potential for future expansion and enhancement.

**Availability and Implementation:** *DrivR-Base* source code is available at https://github.com/amyfrancis97/DrivR-Base.

## 1 Introduction

The rapid advancement of next-generation sequencing technologies has enabled the extensive identification of variants within the human genome, including a substantial proportion classified as variants of uncertain significance. Among these variants, many have the potential to contribute to disease phenotypes, underscoring the critical need to distinguish driver variants from those that are causatively neutral.

In response, a diverse range of machine learning methodologies have been proposed, with the primary objective of using these molecular datasets to identify pathogenic variants. Notable tools in this context include DeepMinds’ most recent piece of work, *AlphaMissense*, (Cheng et al., 2023), our *FATHMM-MKL* (Shihab et al., 2015) and *CScape* (Rogers et al., 2017) predictors, as well as *CADD* (Rentzsch et al., 2019), *DANN* (Quang et al., 2015), *PolyPhen-2* (Adzhubei et al., 2013), and *EVE* (Frazer et al., 2021). While these tools employ diverse methodologies to tackle genomic prediction problems, the datasets, or features, integrated into the models prove equally crucial, and the utility of these classifiers heavily relies on the availability of feature data.

To date, numerous features have demonstrated their effectiveness in assessing the likelihood of a variant driving disease. Conservation-based features, such as PhyloP and PhastCons scores (Pollard et al., 2009; Siepel et al., 2005), quantify sequence conservation across species. Studies have suggested that regions with lower conservation tend to be less functionally significant (Woodruff, 2001). These features have proven informative in several predictors (Sun and Yu, 2019; Cabrera-Alarcon et al., 2022; Shihab et al., 2015; Rentzsch et al., 2019).

Additionally, various other features have played vital roles in driver-variant prediction. For instance, the *Variant Effect Predictor (VEP)* (McLaren et al., 2016) has been instrumental in developing widely-used prediction tools (Shihab et al., 2015; Rentzsch et al., 2019). *VEP* provides valuable insights into variant effects on transcripts within protein-coding regions, introns, and regulatory elements. Moreover, this context has seen the utilization of features such as sequence-based similarity measures, enabling mathematical comparisons of wild-type and mutant string patterns (e.g., spectrum kernels), as well as regulatory features from ENCODE (Dunham et al., 2012; Shihab et al., 2015; Rentzsch et al., 2019; Rogers et al., 2017; Quang et al., 2015). Additionally, information on GC content and CpG islands has proven valuable in these prediction tasks (Shihab et al., 2015; Rogers et al., 2017). Elevated GC content has been associated with increased bendability and the ability to undergo B-Z transitions, which are spatial features linked to open chromatin and active transcription (Vinogradov, 2003).

While various feature groups are currently in use, additional molecular datasets could likely offer valuable insights in predicting driver variants. For instance, exploring the influence of SNVs on DNA shape properties is one such illustration. Multiple DNA shape properties have been implicated in DNA-protein interactions (Jones et al., 2003; Chiu et al., 2017; Rohs et al., 2009). Specifically, high electrostatic potentials have been linked to DNA binding sites (Jones et al., 2003; Chiu et al., 2017), and the narrowing of minor grooves has been associated with A-tracts, resulting in bending towards the minor groove (Rohs et al., 2009). As a result, SNVs occurring at these sites may disrupt these interactions and could lead to functional consequences.

Furthermore, other features that have not been extensively explored in this context include structural information sourced from the AlphaFold (Jumper et al., 2021) and PDB (Berman et al., 2000) databases. These databases contain a wealth of information that could prove valuable when assessing whether a genomic variant is likely to lead to disease. Other examples of feature groups that have not been widely employed thus far and are presented in this work include dinucleotide and amino acid properties.

In this paper, we introduce the creation of a novel repository, named *DrivR-Base*, designed to streamline the data acquisition process for constructing robust predictors of variant driver status. These datasets have broader applications, including the development of haploinsufficiency prediction models (Shihab et al., 2017) and potential adaptation for advancing drug repurposing tools (Irham et al., 2022). We focus on the human genome, providing users with a comprehensive toolkit of scripts, documentation, and links to original sources to build the required feature set. Further details can be found in the Supplementary section.

## 2 Description and Implementation

*DrivR-Base* extracts information for ten different feature groups (FG) from human single nucleotide variants in Browser Extensible Data (BED) format, which are mainly extracted from public databases:

1. **Conservation-based features:** Conservation-based features encompass several crucial metrics. These include PhyloP and PhastCons (Pollard et al., 2009; Siepel et al., 2005) scores, which assess whether nucleotide substitution rates deviate from the expectations under neutral drift. Each of these scores is obtained using seven different alignment methods. Additionally, our analysis incorporates Umap and Bismap mappability data (Karimzadeh et al., 2018), measured using four different types of species alignment methods. These metrics assess the extent to which a genomic region can be accurately mapped during sequencing, providing insights into the reliability of genomic or epigenomic characteristics. Regions exhibiting lower mappability readings may be more prone to error. To obtain these datasets for the entire genome, we retrieve data from the UCSC genome browser (Kent et al., 2002) and tailor our queries to specific input variants. Please note that this script downloads all conservation data locally, temporarily requiring approximately 1TB of disk space.
2. **Variant Effect Predictor:** The *Variant Effect Predictor (VEP)* (McLaren et al., 2016) is organized into three main groups of features. Firstly, we extract all predicted transcript consequences for each variant and encode them using one-hot encoding. The outcome is a file that displays a “1” in the corresponding row for each variant if the transcript consequence is predicted. Next, we retrieve the predicted wild-type and mutant amino acids, presenting the results in two files. The first file follows a BED+2 format, with the final two rows representing the wild-type and mutant amino acids, respectively. For synonymous variants, the amino acids will be the same. Additionally, we generate another file that is one-hot encoded, making it suitable for direct integration into the user’s models. Finally, we extract distances to transcripts. When variants are predicted to affect multiple transcripts, we calculate their mean, maximum, and minimum distances.
3. **Dinucleotide properties:** This feature dataset is sourced from DiProDB, an extensive database encompassing 125 conformational and thermodynamic dinucleotide properties (Friedel et al., 2009), which provides values for four dinucleotide configurations: 1) The wild-type allele paired with the adjacent allele on the left, 2) The wild-type allele paired with the adjacent allele on the right, 3) The mutant allele paired with the adjacent allele on the left, and 4) The mutant allele paired with the adjacent allele on the right. The resulting table contains columns, each representing one of the 125 different properties. Column names include a prefix specifying which of the four configurations it pertains to. For example, ‘1 Propeller Twist’ denotes the value for the propeller twist property in the first configuration.
4. **DNA shape properties:** Here, we incorporate five DNA shape properties from DNAShapeR (Chiu et al., 2016). DNAShapeR employs a sliding-window approach to calculate minor groove width (MGW), helix twist (HelT), propeller twist (ProT), roll (Roll), and electrostatic potential (EP). In our scripts, we extract DNA shape features within a window of +10 and -10 on either side of the variant, but this can be easily adjusted by the user. The output is presented in a table, displaying the value for each DNA shape feature for every calculated base pair, where position 11 corresponds to the variant of interest.
5. **GC content & CpG sites:** *DrivR-Base* also calculates GC content, CpG counts and observed CpG versus expected CpG ratios for nine different window sizes.
6. **Kernel-based sequence similarity:** Our approach also employs sequence-based *p*-spectrum kernels to capture potential disruptions in sequences flanking a single nucleotide variant (Campbell and Ying, 2011). Spectrum kernels allow us to assess the composition of *k*-mers within the genomic regions surrounding a mutation. We explore various window sizes ranging from 2 to 20 and *k*-mer sizes ranging from 1 to 20. For each chosen window size (*w*), we systematically generate all possible combinations of specified *k*-mer sizes for both wild-type and mutant sequences. We then determine the frequency of occurrence for each *k*-mer in the respective sequences using the following mapping function:

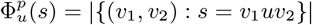

Here, *u* represents the sub-string *k*-mer of length *p, v*1 denotes the wild-type sequence, *v*2 refers to the mutant sequence, and *s* represents the sequence of interest. We subsequently derive a *p*-spectrum kernel by summing the products of corresponding row entries for the two sequences:

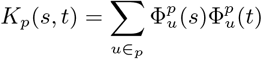

In this equation, *s* corresponds to the wild-type sequence, and *t* corresponds to the mutant sequence. We calculate the diagonals of the *p*-spectra by summing the squares of corresponding row entries within the mapping function matrix. For a more comprehensive explanation and detailed Python implementation, please refer to our Supplementary Material and GitHub Repository.
7. **Amino acid substitution matrices:** In this study, we extract amino acid substitution rates from a variety of matrices for non-synonymous variants sourced from the *Bio*2*mds* package in R (Pelé et al., 2012). The matrices used and their sources are shown in Table 1.
8. **Amino acid properties:** *DrivR-Base* retrieves 532 amino acid properties for both wild-type and mutant amino acid sequences. These properties were sourced from the *AAindex* data within the *AAsea* package in R (Reddy, 2019). They encompass information related to factors such as polarity, hydrophobicity, local flexibility, and helix-bend preferences.
9. **ENCODE database features:** ENCODE offers a wealth of functional information about the human genome (Dunham et al., 2012). In this work, we extract eight features potentially informative for variant pathogenicity:
  i. Transcription Factor ChIP-seq
  ii. Histone ChIP-seq
  iii. DNase-seq
  iv. Mint-ChIP-seq
  v. ATAC-seq
  vi. eCLIP
  vii. ChIA-PET
  viii. GM DNase-seq
  To achieve this, we retrieve all available files for each feature group from ENCODE via the ENCODE API. Subsequently, we download, convert, and consolidate ENCODE peak files into comprehensive data frames for each feature group. These data frames include metadata like accession, target (e.g., transcription factor), biosample (e.g., cell/tissue type), and output type (e.g., narrow peak). Note that this script downloads all ENCODE data locally, requiring approximately 160GB of space. Next, we cross-reference feature-specific databases with target SNVs, extracting relevant information overlapping with SNV locations. We then extract crucial data such as signal values, p-values, q-values, and peaks for each variant. For cases with multiple peaks, such as when replicate assays are involved, we also record minimum, maximum, mean, and range values.
10. **AlphaFold structural features:** *DrivR-Base* incorporates structural data from the AlphaFold database (Jumper et al., 2021) and PDB (Berman et al., 2000). Using the VEP query output, we identify genes and protein positions affected by coding variants. Gene names are converted to UniProtKB IDs, and an API retrieves corresponding crystallographic information files (CIF; .*cif*) from AlphaFold based on the UniProtKB ID. We extract structural information, including X, Y, and Z atom coordinates, isotropic atomic displacement parameters (IADP), and structural conformation types. The output includes two data frames: one containing the first four features (X, Y, Z coordinates, and IADP) for each variant, and another data frame with onehot-encoded structural conformation types indicating potential effects on amino acids, such as bends or helical structures.

**Table 1:**
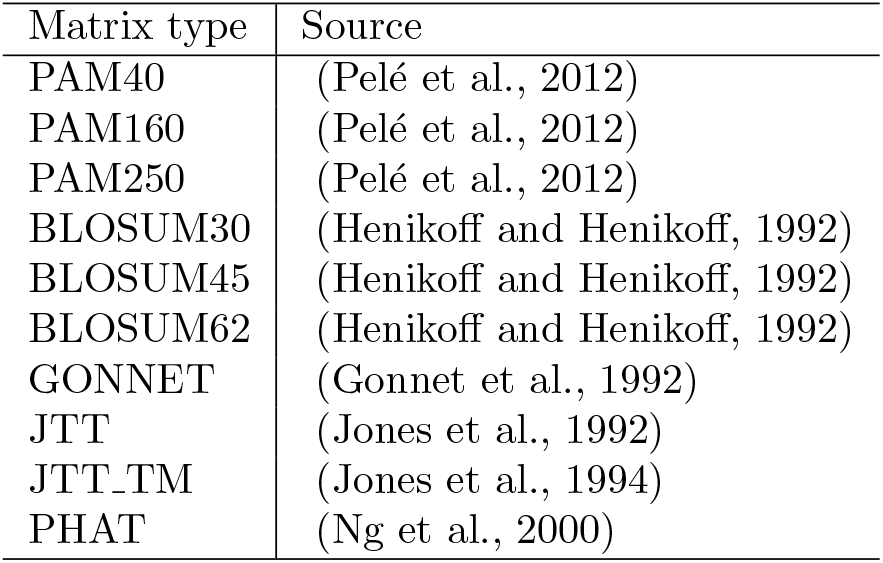
Amino acid substitution matrices and their sources.

A detailed list of feature groups, their sources, and their implementation can be found in our Supplementary Material.

## 3 Conclusions and future efforts

In summary, *DrivR-Base* is a versatile cross-database toolkit that consolidates diverse features for human single nucleotide variants (SNVs). These features have various applications, including constructing high-dimensional machine-learning models for predicting variant driver status. As previously commented, *DrivR-Base* can also be applied to predict haploinsufficient genes and to identify functional similarities to known drug targets, potentially aiding drug repurposing efforts. This tool streamlines feature extraction, saving researchers time and advancing their work. Our future goals include expanding the tool’s capabilities to encompass a broader range of mutations, such as indels, deletions, and structural rearrangements, and diversifying the available feature groups for extraction. Researchers are encouraged to contact the authors to discuss the inclusion of additional feature groups in *DrivR-Base* or the enhancement of existing feature groups.

## 4 Acknowledgements

This work was carried out in the UK Medical Research Council Integrative Epidemiology Unit (MC UU 00032/03) and using the computational facilities of the Advanced Computing Research Centre, University of Bristol.

## 6 Software availablity

DrivR-Base is open sourced and all code is available on GitHub https://github.com/amyfrancis97/DrivR-Base.

