## Supplementary Material for "DrivR-Base: A Feature Extraction Toolkit For Variant Effect Prediction Model Construction"

### DrivR-Base Supplementary Material

January 16, 2024

#### 1 Introduction

In this Supplementary, we present resources available within *DrivR-Base* for downloading data which is informative for predicting the functional impact of variants in the human genome. The scripts in DrivR-Base are organized into molecular feature groups (designated as FG1, FG2, etc.). Within each resource, we provide four subsections to guide the users. Firstly, we provide a brief background explaining why the data source can provide valuable insights for predicting the pathogenic status of a variant. We then provide information about the data sources and locations, and details on how to extract the data, including any useful scripts or tools for the extraction process.
Here, we provide a collection of useful scripts for efficiently aggregating feature data. Our resource compilation, presented below, focuses on the human genome, providing extraction scripts and data location indicators specifically for build GRCh38/hg38 of the human genome.

#### 2 Software Availability

DrivR-Base source code is available at https://github.com/amyfrancis97/DrivR-Base.

#### 3 Feature Groups

##### FG1: Conservation-Based Features

**Background:** Sequence conservation across species serves as a valuable indicator for predicting the pathogenicity of variants. DNA regions which are highly conserved across species have an expectation of being functionally significant. Conversely, there is an indication that variants in poorly conserved regions generally have more limited functional effects and are therefore better tolerated (Woodruff, 2001).

**Data source**: http://hgdownload.cse.ucsc.edu/goldenpath/hg38/

**Script location**: https://github.com/amyfrancis97/DrivR-Base/tree/main/FG1_conservation

**Extraction**: To incorporate conservation-based features, we download 22 datasets comprising Bismap and Umap multi-way mappability scores (Karimzadeh et al., 2018), and PhyloP and PhastCons conservation measures (Pollard et al., 2009; Siepel et al., 2005), from the UCSC Genome Browser Golden Path (Table 1 and 2).

Table 1: Conservation Features

| Conservation Features | |
|---|---|
| phyloP4way | phastCons4way |
| phyloP7way | phastCons7way |
| phyloP17way | phastCons17way |
| phyloP20way | phastCons20way |
| phyloP30way | phastCons30way |
| phyloP100way | phastCons100way |
| phyloP470way | phastCons470way |

Table 2: Mappability Features

| Mappability Features | |
|---|---|
| k24.Bismap.MultiTrackMappability | k24.Umap.MultiTrackMappability |
| k36.Bismap.MultiTrackMappability | k36.Umap.MultiTrackMappability |
| k50.Bismap.MultiTrackMappability | k50.Umap.MultiTrackMappability |
| k100.Bismap.MultiTrackMappability | k100.Umap.MultiTrackMappability |

We then convert the files to bedGraph format using the package: https://anaconda.org/bioconda/ucsc-bigwigtobedgraph. To align the variant file sets with the downloaded datasets, a data reformatting step is necessary to ensure compatibility. Specifically, we transform the original format of chromosome, start, end, reference allele, and alternate allele to the format of chromosome, start-1, end, reference allele, and alternate allele. Once reformatted, we query the downloaded conservation and mappability scores against the variant file using bedtools intersect. The output of this script are 20 separate BED +1 files containing the chromosome location, reference and alternate alleles, and the associated score. Please see the 'get_conservation.sh' script in the '/DrivR-Base/FG1_conservation' folder of our GitHub repository for more details

#### FG2: Variant Effect Predictor

**Background:** Ensembl's Variant Effect Predictor (VEP) (McLaren et al., 2016) is a tool that offers a diverse array of features that can be predicted for single nucleotide variants. The features used in this analysis include predicted consequences on transcripts, distances from transcripts, and amino acids that are predicted to be impacted by the variant. The consequences that are predicted by VEP and a description can be found in Table 3. Predicting effects on transcripts offers valuable insights into functional significance. For example, variants that are predicted to lead to more severe consequences, such as the introduction of premature stop codons, or frameshift variants, are more likely to be pathogenic and to play a role in disease. The predicted amino acids from VEP enable us to establish whether a variant is synonymous and whether a non-synonymous variant leads to an amino acid whose chemical properties differ substantially from the wild type. Hence, by considering the predicted amino acid changes, we gain insights into the potential functional implications of variants and their association with disease.

Table 3: Consequences of Genetic Variants

| Consequence | Description |
|---|---|
| splice_acceptor_variant | Variant affects splice acceptor site |
| splice_donor_variant | Variant affects splice donor site |
| stop_gained | Gain of a stop codon |
| frameshift_variant | Indel causes a frameshift |
| stop_lost | Loss of a stop codon |
| start_lost | Loss of a start codon |
| transcript_amplification | Amplification of a transcript |
| feature_elongation | Elongation of a genomic feature |
| feature_truncation | Truncation of a genomic feature |
| inframe_insertion | In-frame insertion of bases |
| inframe_deletion | In-frame deletion of bases |
| missense_variant | Missense mutation |
| protein_altering_variant | Protein-altering variant |
| splice_donor_5th_base_variant | Variant at the 5th base of a splice donor site |
| splice_region_variant | Variant in a splice region |
| splice_donor_region_variant | Variant in a splice donor region |
| splice_polypyrimidine_tract_variant | Variant in a polypyrimidine tract |
| incomplete_terminal_codon_variant | Incomplete terminal codon |
| start_retained_variant | Start codon retained |
| stop_retained_variant | Stop codon retained |
| synonymous_variant | Synonymous mutation |
| coding_sequence_variant | Variant in coding sequence |
| mature_miRNA_variant | Variant in mature miRNA |
| 5_prime_UTR_variant | Variant in 5' untranslated region |
| 3_prime_UTR_variant | Variant in 3' untranslated region |
| non_coding_transcript_exon_variant | Variant in non-coding transcript exon |
| intron_variant | Variant in intron |
| NMD_transcript_variant | NMD (Nonsense-Mediated Decay) transcript variant |
| non_coding_transcript_variant | Variant in non-coding transcript |
| coding_transcript_variant | Variant in coding transcript |
| upstream_gene_variant | Variant upstream of a gene |
| downstream_gene_variant | Variant downstream of a gene |
| TFBS_ablation | Transcription factor binding site (TFBS) ablation |
| TFBS_amplification | Transcription factor binding site (TFBS) amplification |
| TF_binding_site_variant | Variant affects TF binding site |
| regulatory_region_ablation | Regulatory region ablation |
| regulatory_region_amplification | Regulatory region amplification |
| regulatory_region_variant | Variant affects regulatory region |
| intergenic_variant | Intergenic variant |
| sequence_variant | Generic sequence variant |

**Data source**: https://www.ensembl.org/info/docs/tools/vep/script/vep_cache.htmlcache

**Script location**: https://github.com/amyfrancis97/DrivR-Base/tree/main/FG2_vep

**Extraction**: To utilize the Variant Effect Predictor (VEP) for variant annotation, we begin by downloading the VEP cache to our local environment. A list of prerequisites and dependencies for cache download can be found in our DrivR-Base GitHub repository. Additional information is available in the VEP cache tutorial online. Once the VEP cache is downloaded locally, we can proceed to query and extract relevant features for specific variants. For a detailed description of the querying process, please refer to our 'get_vep.sh' script located in the '/DrivR-Base/FG2_vep' sub-directory of the repository.

We extract these features as separate queries to make the output more interpretable and, after querying features for the given variant, we reformat this output to similarly enhance useability. For consequence features, we implement a one-hot-encoding approach. Each possible consequence is represented as a distinct column. A '1' in a column indicates the prediction of a specific consequence, while '0' denotes its absence. Similarly, for amino acid features, we concatenate two one-hot-encoded vectors. The first vector corresponds to the wild-type amino acid, with each element representing one of the 20 amino acids. A '1' in the vector indicates the amino acid coded by the wild-type codon. The second vector follows the same structure but relates to the mutant amino acid. The resulting two vectors are then concatenated to form the final amino acid feature representation.

#### FG3: Dinucleotide Properties

**Background:** In our study, we use features derived from DiProDB (Friedel et al., 2009), a comprehensive database of conformational and thermodynamic dinucleotide properties. We can use this information to elucidate any conformational and thermodynamic changes that occur between wild-type and mutant genomic sequences, hence providing insight into the potential functional significance of variants on a DNA-level.

**Data source**: https://diprodb.fli-leibniz.de/ShowTable.php

**Script location**: https://github.com/amyfrancis97/DrivR-Base/tree/main/FG3_dinucleotide_properties

**Extraction**: For each variant, we extract values for four dinucleotide configurations in R: 1) The wild-type allele paired with the adjacent allele on the left, 2) The wild-type allele paired with the adjacent allele on the right, 3) The mutant allele paired with the adjacent allele on the left, and 4) The mutant allele paired with the adjacent allele on the right. Each configuration contains 125 dinucleotide property features, resulting in a dataframe containing 500 feature columns for each variant, with corresponding row values from the original DiProDB table. The script for this extraction process is 'dinucleotide_properties.R' and can be found in '/DrivR-Base/FG3_dinucleotide_properties'

#### FG4: DNA Shape Properties

**Background:** We use DNAShapeR (Chiu et al., 2016) to capture DNA shape properties of genomic variant sites. Specifically, this package uses a sliding pentamer window approach and Monte Carlo simulations to determine the minor groove width (MGW), helix twist (HelT), propeller twist (ProT), and roll (Roll) values for DNA sequences. Crucially, DNA shape features are well-established for their roles in protein-DNA binding specificity (Jones et al., 2003; Chiu et al., 2017; Rohs et al., 2009). Given the significance of these DNA shape features in protein-DNA interactions, it is reasonable to hypothesize that variants that impact these shapes may have greater pathogenic potential. By incorporating these DNA shape features into our analysis, we aim to capture the impact of variants on protein-DNA binding and explore their potential association with pathogenicity.

**Data source**: https://www.ncbi.nlm.nih.gov/pmc/articles/PMC4824130/

**Script location**: https://github.com/amyfrancis97/DrivR-Base/tree/main/FG4_dna_shape

**Extraction**: We read variant datasets into R and extract DNA nucleotides flanking ten base pairs either side of the variant of interest. This results in a 21 base-pair DNA sequence, where base 11 denotes the variant. We then pass this sequence to the 'getShape()' function in 'DNAShapeR'. This calculates MGW, HelT, ProT, and Roll values for each of the positions in the wild-type nucleotide sequence. This enables us to establish whether variants found in regions exhibiting particular dna shape properties are more likely to lead to pathogenic effects. For detailed steps on the implementation of this process, please refer to the 'dna_shape.R' script located in the '/DrivR-Base/FG4_dna_shape' directory of the repository.

#### FG5: GC and CpG content

**Background:** The GC content, also referred to as the *isochore* structure, exhibits significant heterogeneity in the mammalian genome. While many questions still remain regarding the functional implications of these organizational events, studies have associated these structures with various genomic features that hold potential functional significance in the context of driver prediction. For example, increased bendability and B-Z transitions have been associated with elevated GC content and, these structures have been linked to open chromatin and active transcription (Vinogradov, 2003). DNA curvature has been associated with condensed chromatin state (Vinogradov, 2003). This feature group also include CpG counts and observed vs expected ratios. CpG sites are frequently enriched in gene promoters within mammalian genomes (Rozenberg et al., 2008). Hence, there is potentially a gain to be made in including these feature groups when predicting the functional impact of genomic variants.

**Script location**: https://github.com/amyfrancis97/DrivR-Base/tree/main/FG5_gc_CpG

**Extraction**: We first download the GRCh38 human genome fasta files from NCBI (Sayers et al., 2022). We then query the genome to extract nucleotide sequences surrounding the variant of interest. We repeat this for 9 different window sizes. For each of these sequences, we then calculate GC content:

$$GCcontent = \frac{Count(G + C)}{Count(A + T + G + C)} \times 100\% \qquad (1)$$

Next, we calculate the observed vs expected ratio:

$$Obs/Exp(CpG) = \frac{Count(CpG)}{Count(C + G)} \times n \qquad (2)$$

Where $Count(CpG)$ is the number of CG dinucleotides, and $n$ is the nucleotide sequence length.

#### FG6: Kernel-Based Sequence Similarity

**Background:** We employ sequence-based $p$-spectrum kernels to capture potential disruptions in sequences flanking a single nucleotide variant (Campbell and Ying, 2011). Spectrum kernels are used to characterize the composition of $k$-mers within regions surrounding the SNV, both before and after the mutation occurs. The result of the spectrum kernel algorithm is the generation of two $k$-spectra: one representing the wild-type version of the sequence and the other representing the mutant version. These $k$-spectra encode the frequency and distribution of $k$-mers within their respective sequences. By concatenating these two $k$-spectra, we obtain a comprehensive representation that encapsulates the complete picture of how the sequence changes due to the SNV.

**Script location**: https://github.com/amyfrancis97/DrivR-Base/tree/main/FG6_kernel

**Extraction**: Similarly to the previous feature group, we first download the GRCh38 human genome fasta files from NCBI (Sayers et al., 2022). We then query the genome to extract nucleotide sequences surrounding the variant of interest for window sizes ranging between 2 and 10. In the 'get_kernel.py' script, we created an 'over_slice()' function which systematically generates all possible combinations of $k$-mers of given sizes, $k$, for each pair of wild type and mutant sequences using a sliding window approach. The local sequence patterns around the variants can then be calculated from these $k$-mers. To count the occurrence of each $k$-mer, we employ a mapping function:

$$\Phi_u^p(s) = |\{(v_1, v_2) : s = v_1 u v_2\}|$$

Here, $u$ represents the sub-string $k$-mer of length $p$, $v_1$ denotes the wild-type sequence, $v_2$ refers to the mutant sequence, and $s$ represents the sequence of interest. The mapping function quantifies the number of times the substring $u$ appears in the sequence $s$, considering both the wild-type and mutant sequences. For example, let's consider the sequences "ATCGT" and "ATAGT", where the mutant sequence has replaced a cytosine residue with an adenine. If we use a $k$-mer size of 2, the result of the mapping function would be:

Table 4: p-spectrum kernel mapping function

| **seq** | **AT** | **TC** | **CG** | **GT** | **TA** | **AG** |
|---|---|---|---|---|---|---|
| ATCGT (s) | 1 | 1 | 1 | 1 | 0 | 0 |
| ATAGT (t) | 1 | 0 | 0 | 1 | 1 | 1 |

From the mapping function, we can derive a $p$-spectrum kernel matrix by summing the products of the corresponding row entries for the two sequences:

$$K_p(s,t) = \sum_{u \in_p} \Phi_u^p(s) \Phi_u^p(t)$$

In this equation, $s$ corresponds to the wild-type sequence, and $t$ corresponds to the mutant sequence. The practical procedure involves the following steps:

1) Count the number of 'AT' $k$-mers in the wild-type sequence ($s$), which results in a count of 1.
2) Count the number of 'AT' $k$-mers in the mutant sequence ($t$), which also results in a count of 1.
3) Multiply the counts obtained in steps 1 and 2 (1 * 1 = 1) for each of the 6 $k$-mer types.
4) Sum the results from step 3 to obtain the final value.

We calculate the diagonals of the $p$-spectra by summing the squares of corresponding row entries within the mapping function matrix for both sequences $s$ and $t$, using the equation:

$$K_p(z, z) = \sum_{u \in_p} (\Phi_u^p(z))^2$$

In this equation, $z$ represents either $s$ or $t$, and we apply this calculation to both sequences to determine the diagonals for each.

For our example, we calculate this in Python using the following steps:

1) For the wild-type sequence ($s$), we square the row entry for each of the 6 $k$-mers, and then sum the result. This results in a count of 4 as each row entry is 1.
2) For the mutant sequence ($t$), we repeat the above process. This also results in a count of 4.

The resulting $p$-spectrum kernel matrix is:

Table 5: p-spectrum kernel matrix

| **K** | **ATCGT (s)** | **ATAGT (t)** |
|---|---|---|
| **ATCGT (s)** | 4 | 2 |
| **ATAGT (t)** | 2 | 4 |

For a more comprehensive explanation and detailed Python implementation, please see the 'get_kernel.py' script in our DrivR-Base GitHub Repository.

#### FG7: Amino Acid Substitution Matrices

**Background:** Amino acid substitution matrices play a crucial role in understanding the evolutionary dynamics of protein sequences. These matrices quantify the rates at which amino acid residues are substituted for one another during evolution. They provide valuable information about the likelihood of specific amino acid substitutions occurring in proteins. Amino acids that have a low probability of being substituted are typically those that differ significantly in their chemical composition, structure, or function. These substitutions are less common due to the functional and structural constraints imposed on protein sequences. Therefore, when a variant leads to an amino acid substitution that is associated with a low substitution probability according to the amino acid substitution matrix, it suggests a substantial change in the chemical composition of the protein. Such substitutions may have a greater likelihood of impacting the protein's function, stability, or interaction with other molecules.
By considering amino acid substitution matrices, we can gain insights into the potential functional consequences of amino acid changes caused by variants. For variants in coding regions of the genome, this information is valuable for predicting pathogenicity and understanding their

impacts on protein structure and function.

**Data source**: https://rdrr.io/cran/bios2mds/man/sub.mat.html

**Script location**: https://github.com/amyfrancis97/DrivR-Base/tree/main/FG7_aa_substitution_matrices

**Extraction**: To incorporate the amino acid substitution information, we use predicted wild-type and mutant amino acids obtained from the Variant Effect Predictor (VEP) output. We compared these amino acids against a collection of eight widely used amino acid substitution matrices, which were sourced from the 'sub.mat' data in the 'bio2mds' R package. Table 6 shows the substitution matrices used and their sources.

Table 6: Amino acid substitution matrices and their sources

| Substitution Matrix | Source |
|---|---|
| PAM40 | (Pelé et al., 2012) |
| PAM160 | (Pelé et al., 2012) |
| PAM250 | (Pelé et al., 2012) |
| BLOSUM30 | (Henikoff and Henikoff, 1992) |
| BLOSUM45 | (Henikoff and Henikoff, 1992) |
| BLOSUM62 | (Henikoff and Henikoff, 1992) |
| GONNET | (Gonnet et al., 1992) |
| JTT | (Jones et al., 1992) |
| JTT_TM | (Jones et al., 1994) |
| PHAT | (Ng et al., 2000) |

#### FG8: Amino Acid Properties

**Background:** To capture structural information and functional impacts of amino acid changes, we incorporate properties for the 20 naturally occurring amino acids using the 'AAindex' list from the 'AAsea' package in R (Reddy, 2019). The 'AAindex' list is a comprehensive collection of 536 amino acid properties compiled from various reliable sources and contains a range of important features, such as *Cruciani* properties (principal components representing amino acid interactions with chemical groups), physical properties (including hydrophobicity, helix bends, and side chain size), and chemical properties (including pKa values and chemical shifts).
Hydrophobicity, for example, plays a role in protein folding and stability, impacting the organisation and packaging of protein structures. The Cruciani properties capture the interactions between amino acids and specific chemical groups, providing information about their chemical behaviour and potential functional significance. The physical properties, including helix bends and side chain size, contribute to the three-dimensional structure and function of proteins. Additionally, pKa values provide insights into the ionization behaviour and acid-base properties of amino acids.
By incorporating these amino acid properties as encoders for the impacted amino acids in each variant, we aim to capture functionally significant physical and chemical properties between wild-type and mutant amino acids.

**Data source**: https://github.com/cran/aaSEA/tree/master/data

**Script location**: https://github.com/amyfrancis97/DrivR-Base/tree/main/FG8_aa_properties

**Extraction**: We use the predicted amino acid output from our FG2_vep feature group as input for these scripts. To execute these scripts, we must first ensure that the 'AAindex.rda' R data file is downloaded and that the 'config.R' file is updated with the location in which this data file has been stored. We have provided the R data file in the '/DrivR-Base/FG8_aa_properties' sub-folder. We load this dataset into our R environment and query the values for each of the 536 properties, repeating this step for both the wild-type and mutant amino acids. We then concatenate these two dataframes by column. The resulting file contains the values for each of the wild-type and mutant amino acids for every variant predicted to be coding.

#### FG9: ENCODE Regulatory Features

**Background:** The ENCODE consortium is a comprehensive database that provides valuable information on regulatory regions within the DNA (Dunham et al., 2012). Through various experimental approaches, such as investigating open chromatin states, histone enrichment, transcription factor binding, gene expression, RNA binding sites, and methylation, ENCODE offers a wealth of data that can shed light on the functional activity of genomic sites.
While these functionally related elements can play roles in coding regions of the DNA, such as the enrichment of acetylated histones, their significance is often more pronounced in non-coding regions. These functional elements play crucial roles in the regulation of transcription and gene expression. Open chromatin regions, for instance, have been found to interact with regulatory elements, while transcription factors bind to promoters or enhancers to tightly regulate gene expression (Hu and Tee, 2017). As a result, variants located within regions enriched with these regulatory elements have a higher likelihood of having functional impacts.
ENCODE serves as a valuable resource for identifying and understanding these functional elements, providing insights into the regulatory mechanisms that govern gene expression and transcriptional regulation. By leveraging the data available in ENCODE, we can gain a deeper understanding of the functional significance of genomic variants and their potential impact on gene regulation and cellular processes.

**Data source**: https://www.encodeproject.org/

**Script location**: https://github.com/amyfrancis97/DrivR-Base/tree/main/FG9_encode

**Extraction**: First, we download all available ENCODE files for eight different features (shown in Table 7) and convert them from bigBed to BED format.

Table 7: ENCODE Features

| **ENCODE Features** | |
|---|---|
| TF ChIP-seq | Histone ChIP-seq |
| DNase-seq | Mint-ChIP-seq |
| ATAC-seq | eCLIP |
| ChIA-PET | GM DNase-seq |

Subsequently, we concatenate all the separate files into a single file per feature group, from which we can later extract features by querying variants. We have provided the 'downloadEncode.py' script in the 'DrivR-Base/FG9_encode' directory and have also included the downloaded datasets as zipped files in the supplementary data. Please note that this script downloads all ENCODE data locally, requiring approximately 160GB of disk space.
We then extract signal values, p-values, q-values, and peaks for each unique biosample and target. For instance, Transcription Factor ChIP-Seq involves multiple transcription factor targets (e.g., CTCF) measured in various tissues (e.g., BLaER1). In cases where there are multiple values for a given biosample and target, such as replicate assays in a specific tissue, we calculate and record the mean, minimum, maximum, and range. The resulting file contains the variant location, followed by individual values for each unique biosample and target (e.g., CTCF_BLaER1_mean).

#### FG10: Alpha Fold Structural Features

**Background:** Alphafold and PDB, used together, have enabled the prediction of a vast number of protein structures (Jumper et al., 2021; Berman et al., 2000). Once we identify the protein and the specific position at which a non-synonymous coding variant occurs, we can utilize this data to enhance our understanding of the chemical and physical composition of the surrounding protein region. In this context, we extract five atomic features and two structural conformation features from the AlphaFold crystallographic information files (CIF; *.cif*). Specifically, we retrieve the x, y, and z coordinates, which are defined with respect to a set of orthogonal Cartesian axes. This description pertains to the atomic coordinates found in crystallographic structural data and is presented in Table 8.

**Data source**: https://alphafold.ebi.ac.uk/

**Script location**: https://github.com/amyfrancis97/DrivR-Base/tree/main/FG10_alpha_fold

**Extraction**: We use the full VEP query output from our FG2_vep feature group as input for these scripts ('file_variant_effect_output_all.txt'). The 'get_alpha_fold.py' script extracts the predicted gene names and predicted protein positions from the VEP file and converts the gene names to UniProtKB IDs. We then use the 'requests' library to fetch the appropriate file for a given variant and read it into Python. Next, we extract the relevant values for each of the structural and atom-site features at the specified protein positions. For the structural conformation features, we convert the output into a one-hot-encoded table, where a '1' in the conformation type column represents the presence of that type.

Table 8: Alpha Fold Features

| Feature | Label | Description |
|---|---|---|
| X Atom-Site Coordinate | _atom_site.Cartn_x | The x atom-site coordinate in angstroms specified according to a set of orthogonal Cartesian axes related to the cell axes |
| Y Atom-Site Coordinate | _atom_site.Cartn_y | The y atom-site coordinate in angstroms specified according to a set of orthogonal Cartesian axes related to the cell axes |
| Z Atom-Site Coordinate | _atom_site.Cartn_z | The z atom-site coordinate in angstroms specified according to a set of orthogonal Cartesian axes related to the cell axes |
| Atom-Site Occupancy | _atom_site.occupancy | The fraction of the atom type present at this site |
| Isotropic Displacement Parameter | _atom_site.B_iso_or_equiv | Isotropic atomic displacement parameter, or equivalent isotropic atomic displacement parameter, Beq, calculated from the anisotropic displacement parameters |
| Structural Conformation Type | _struct_conf.conf_type_id | The type of structural conformation (e.g., BEND) |
| Structural Conformation ID | _struc_conf.id | The ID of the structural conformation (e.g., BEND5) |

Source: The feature descriptions presented in this table have been taken or adapted from the PDBx/mmCIF dictionary (https://mmcif.wwpdb.org/pdbx-mmcif-home-page.html).
